# Potentiation of glutamatergic synaptic transmission onto lateral habenula neurons following early life stress and intravenous morphine self-administration in rats

**DOI:** 10.1101/2020.12.23.424217

**Authors:** Ludovic D. Langlois, Rina Y. Berman, Ryan D. Shepard, Sarah C. Simmons, Mumeko C. Tsuda, Shawn Gouty, Kwang H. Choi, Fereshteh S. Nugent

## Abstract

Early life stress (ELS) presents an important risk factor for drug addiction and comorbid depression and anxiety through persistent effects on the mesolimbic dopamine (DA) pathways^1^. Using an ELS model for child neglect (a single 24 h episode of maternal deprivation, MD) in rats, recent published works from our lab show that MD induces dysfunction in ventral tegmental area (VTA) DA neurons ^2–4^ and its negative controller, the lateral habenula (LHb) ^5–7^. In regard to LHb, MD-induced potentiation of glutamatergic synaptic transmission onto LHb neurons shifts the coordination of excitation/inhibition (E/I) balance towards excitation, resulting in an increase in the overall spontaneous neuronal activity with elevation in bursting and tonic firing, and intrinsic excitability of LHb neurons in early adolescent male rats ^5–7^. Here, we explored how MD affects intravenous morphine self-administration (MSA) acquisition and sucrose preference as well as glutamatergic synaptic function in LHb neurons of adult male rats self-administering morphine. We found that MD-induced increases in LHb neuronal and glutamatergic synaptic activity and E/I ratio persisted into adulthood. Moreover, MD significantly reduced morphine intake, triggered anhedonia-like behavior in the sucrose preference test (SPT), and was associated with persistent glutamatergic potentiation 24h after the last MSA session. MSA also triggered postsynaptic glutamatergic potentiation in LHb neurons of control rats during this time period. Our data highlights that ELS-induced glutamatergic plasticity in LHb may dampen the positive reinforcing properties of natural rewards and opioids, and contribute to the development of anhedonic and dysphoric states associated with opioids.

## Introduction

Exposure to early life stress (ELS) is a strong predictor for several later-life mental disorders, including substance use disorders, anxiety and depression ^8–11^. ELS may increase the risk of a variety of mental illnesses and addiction through modifications in brain reward circuits and synaptic integration controlling brain dopamine (DA) signaling ^1,3,6,12–15^ although the exact mechanistic link between ELS and this increased vulnerability is still unknown.

Prevalent ELS rodent models are maternal deprivation (MD, a single prolonged episode of maternal deprivation) and maternal separation (MS, repeated daily maternal separations) models, in which animals are separated from their mothers early in life. Our recent data using an established rodent ELS model of child neglect in Sprague Dawley (SD) rats with a single 24 h episode of MD at postnatal day 9 (PND9) indicates that the lateral habenula (LHb) may serve as a critical converging brain region for ELS-induced dysregulation of reward circuits including ventral tegmental area (VTA) DA signaling ^5–7^.

The LHb is an emerging anti-reward hub for motivation and decision making that links forebrain limbic structures with midbrain monoaminergic centers ^16–19^ and is involved in reward/aversion-related learning and memory processing associated with avoidance from stressful and aversive situations through the suppression of DA and serotonin systems. Not surprisingly, LHb dysfunction contributes to a myriad of cognitive, learning, and affective impairments associated with depression, anxiety, psychosis, and drug addiction ^20–22^. Consistently, we also found that MD produces pro-depressive-like behaviors in the forced swim test (FST) in late adolescent male rats, increases spontaneous neuronal activity (increased bursting and tonic firing patterns) and intrinsic excitability of LHb neurons in early and late adolescent male rats ^5–7^. MD is also associated with pre- and post-synaptic potentiation of AMPA receptor (AMPAR)-mediated glutamatergic transmission that shifts excitation/inhibition (E/I) balance towards excitation supporting LHb hyperactivity ^5,6^, although it remains unknown if MD-induced neuroadaptations and LHb hyperactivity persist into adulthood. Here, we first investigated whether early MD stress induces persistent LHb neuronal and synaptic dysfunction in adult male rats. Moreover, we investigated the effects of MD on sucrose preference and intravenous morphine self-administration (MSA) acquisition behavior given that the effects of ELS on opioid intake are conflicting, and the interaction between ELS and the perception of reward is a subject of ongoing research. ELS research typically focuses on intake patterns of stimulants and alcohol, but overlooks opioids, including morphine, which are particularly addictive. A few reports have shown that MS (using a 4 h daily isolation of pups from litters from PND1-PND14) induces a higher dependence on morphine in the conditioned place preference test (CPP with intraperitoneal drug administration) and oral self-administration models in Long–Evans adult male rats ^23–25^. Mu opioid receptor (MOR) activation is found to suppress LHb activity ^26^ and naloxone-precipitated morphine withdrawal from passive and repeated injections of morphine in mice depresses glutamatergic synapses onto raphe-projecting LHb neurons ^27^. Nevertheless, the effects of intravenous MSA on LHb glutamatergic function and following MD are still unknown. Here, we demonstrate that MD alters MSA acquisition and decreases sucrose preference, and is associated with persistent MD-induced LHb glutamatergic synaptic plasticity that supports enhanced E/I ratio and LHb hyperactivity. This persistent MD-induced glutamatergic potentiation in LHb neurons may contribute to anhedonia to natural rewarding stimuli (such as sucrose in the SPT) and opioids (morphine in the MSA) in adult male SD rats.

## Materials and methods

### Animals

All experiments employed male SD rats (sourced from Taconic Inc, and Charles River Laboratories) in experiments conducted in accordance with the National Institutes of Health Guide for the Care and Use of Laboratory Animals and were approved by the Uniformed Services University Institutional Animal Care and Use Committee. All rats were received on PND6 with lactating dams and allowed to acclimate undisturbed for ~72h before initiation of the MD procedure. All rats were kept on a 12h dark: 12h light cycle schedule with lights on at 06:00, and all procedures began 3–4 h after the start of the light-cycle (except for SSA and MSA procedure, see below). All animals received ad libitum standard chow and water (except where noted during the MD procedure).

### MD procedure

MD was performed on male rats at PND9. Half of the rats in the litter (randomly selected) were isolated from the dam and their siblings for 24h (MD group). The isolated pups were placed on a heating pad (34°C) in a separate quiet room and not disturbed for 24h until being returned to their home cage. The remaining non-maternally deprived control group (non-MD) received the same amount of handling as the MD rats. All rats were group-housed (2 per cage, treatment-matched) from weaning at PND28 until sacrifice at the age range of PND70-PND80 with standard housing care and no additional experimenter manipulation prior to sacrifice.

### Slice Preparation

For all electrophysiology experiments, several separate cohorts of non-MD/MD-treated rats were used. As described before ^6^, all rats were anesthetized with isoflurane, decapitated, and brains were quickly dissected and placed into ice-cold artificial cerebrospinal fluid (ACSF) containing (in mM): 126 NaCl, 21.4 NaHCO_3_, 2.5 KCl, 1.2 NaH_2_PO_4_, 2.4 CaCl_2_, 1.0 MgSO_4_, 11.1 glucose, and 0.4 ascorbic acid and saturated with 95% O_2_-5% CO_2_. Briefly, sagittal slices containing LHb were cut at 250 μm and incubated in above prepared ACSF at 34°C for at least 1 h prior to electrophysiological experiments. For patch clamp recordings, slices were then transferred to a recording chamber and perfused with ascorbic-acid free ACSF at 28°C.

### Electrophysiology

Voltage-clamp cell-attached and voltage/current-clamp whole-cell recordings were performed from LHb neurons in LHb-containing slices using patch pipettes (3-6 MOhms) and a patch amplifier (MultiClamp 700B) under infrared-differential interference contrast microscopy. Data acquisition and analysis were carried out using DigiData 1440A, pCLAMP 10 (Molecular Devices), Clampfit, and Mini Analysis 6.0.3 (Synaptosoft, Inc.). Signals were filtered at 3 kHz and digitized at 10 kHz.

To assess LHb spontaneous activity, cells were patch clamped with potassium gluconate-based internal solution (130 mM K-gluconate, 15 mM KCl, 4 mM adenosine triphosphate (ATP)-Na^+^, 0.3 mM guanosine triphosphate (GTP)-Na^+^, 1 mM EGTA, and 5 mM HEPES, pH adjusted to 7.28 with KOH, osmolarity adjusted to 275-280 mOsm) in slices perfused with ACSF. Spontaneous neuronal activity and AP firing patterns (tonic, bursting) were assessed in both cell-attached recordings in voltage-clamp mode at V=0 and whole-cell recording in current-clamp mode at I=0 for the duration of the ~1 min recording. Cells that fired less than 2 action potentials (APs) during this time period were characterized as silent.

The excitatory and inhibitory balance (E/I ratio) was recorded with a cesium (Cs)-gluconate– based internal solution in intact synaptic transmission. Patch pipettes were filled with Cs-gluconate internal solution (117 mM Cs-gluconate, 2.8 mM NaCl, 5 mM MgCl_2,_ 2 mM ATP-Na^+^, 0.3 mM GTP-Na^+^, 0.6 mM EGTA, and 20 mM HEPES, pH adjusted to 7.28 using CsOH, osmolality adjusted to 275-280 mOsm). Evoked EPSCs and IPSCs from LHb neurons were recorded in the same neuron in drug-free ACSF using a stimulating electrode placed in the stria medullaris. EPSCs were recorded at the reversal potential for GABA_A_ IPSCs (−55 mV), and IPSCs were recorded at the reversal potential for EPSCs (+10 mV) in the same LHb neuron. The E/I ratio was then calculated as EPSC/IPSC amplitude ratio by dividing the average peak amplitude of 10 consecutive sweeps of EPSCs or IPSCs from the same recording. Whole cell recordings of AMPAR-meditated miniature excitatory postsynaptic current (mEPSC) recordings were performed in ACSF perfused with picrotoxin (100 μM), d-APV (50 μM), and tetrodotoxin (TTX, 1 μM), and pipettes filled with Cs-gluconate internal solution similar to E/I ratio recordings. For mEPSCs, LHb neurons were voltage-clamped at −70 mV and recorded over 10 sweeps, each lasting 50 s. The cell series resistance was monitored through all the experiments and if this value changed by more than 10%, data were not included.

### Catheter Surgery

Non-MD and MD male rats (PND60-PND63) were anesthetized with a cocktail of ketamine/xylazine (100 mg/kg and 10 mg/kg, i.p.) and a catheter was implanted in the right jugular vein and attached to the back of the animal as described previously ^28,29^. The entry point of the catheter was secured in place with a 0.5 cm × 0.5 cm Mersilene surgical mesh attached to the catheter. Then the catheter composed of 22-gauge stainless steel tubing cemented into place with bell-shaped dental cement with a 1.5cm × 1.5 cm Marlex surgical mesh was secured under the skin of the back of the animal. The incision was closed using stainless steel wound clips and treated with topical antibiotic ointment. Animals received antibiotic gentamycin sulfate (5-8 mg/kg, i.v.) after the surgery and were monitored until full recovery.

### Morphine self-administration (MSA)

Eight operant conditioning chambers (Med Associates Inc., St. Albans, VT) were used for intravenous MSA experiments. Each chamber was equipped with two levers, an infusion pump, and a 10 mL glass syringe connected to a fluid swivel (Instech, Plymouth Meeting, PA) by Teflon tubing. After one week recovery from catheter surgery, animals (PND67-PND70) were placed in the operant conditioning chambers and allowed to self-administer either morphine (0.3 mg/kg/infusion, 0.1 mL over 5 sec, MSA) or 0.9% saline (saline self-administration, SSA) on one lever press/injection (Fixed Ratio, FR 1) reinforcement schedule for a 4 h session with a maximum number of infusions 60 a day. Animals were trained with a FR1 schedule of reinforcement for 6 days, and then switched to a FR3 schedule from days 7 to 10. During each infusion (5 sec), a cue light above the active lever was illuminated, and the house light was extinguished for an additional 20 sec (time-out) period after infusion. The lever press responses on the inactive lever and responses to both levers during the time-out period were recorded but had no programmed consequences. Each chamber was equipped with two infrared photo beams that monitored spontaneous locomotor activity of animals during the self-administration session. The numbers of drug-paired lever presses, drug infusions, and inactive lever presses were recorded by the MED-PC®-IV self-administration software.

### Sucrose Preference Test (SPT)

For the SPT, experiments were conducted when animals were transferred to a reverse- light cycle. At PND50, non-MD and MD male rats were transferred to a reverse light cycle room (12:12, light: dark, lights off at 0600) and acclimated to the new light cycle for one week. At PND58, non-MD and MD rats began habituation to drinking a 2% sucrose solution in their home cage over a one-week period. During habituation, rats were presented with a two-bottle choice selection containing either a 2% sucrose solution or water. Each habituation day, water and sucrose bottles positions were alternated and checked for leakage. Prior to testing, both bottles were removed from the cage and animals underwent 18 h of water deprivation. At PND70, animals were individually placed in a new cage allowed 30 minutes to acclimate to the new cage, and then presented with the two-bottle choice selection for a one-hour test period. After the test, bottles were removed and weighed. Sucrose Preference (% Preference) was calculated using the following formula: % Preference = [(sucrose consumed in g)/(sucrose consumed in g+ water consumed in g)] × 100.

### Statistics

Values are presented as mean ± SEM. The threshold for significance was set at *p < 0.05 for all analyses. All statistical analyses of data were performed using GraphPad Prism 8.4.1. For the effects of MD on E/I ratios and the SPT, we used unpaired Student’s t test. For detecting the difference in distribution of silent, tonic or bursting LHb neurons in non-MD and MD rats, we used Chi-square tests. Mini Analysis software was used to detect and measure mEPSCs using preset detection parameters of mEPSCs with an amplitude cutoff of 5 pA. Effects of MD and MSA on the mean and cumulative probabilities of mEPSC amplitude and frequency data sets were analyzed using two-way ANOVA tests and Kolmogorov-Smirnov (KS) tests, respectively. Effects of MD stress and MSA during the acquisition time were analyzed separately for the FR1 and FR3 schedules using 3-way Repeated Measures (RM) ANOVA and Mixed-effect (only for locomotor activity with the FR1 schedule) tests for with Tukey’s post-hoc test with MD and MSA as between-group factors and time as the within-group factor.

## Results

### MD-induced changes in LHb E/I balance and spontaneous activity persisted into adulthood

Here, we first examined whether MD-induced increases in E/I ratio and LHb neuronal activity persist in adult male rats. We detected an increase in E/I ratio following MD suggesting a shift in the balance between excitation and inhibition in synaptic inputs towards excitation following MD (Figure 1A, t (31) = 2.07, p < 0.05). We observed that MD increased LHb spontaneous neuronal activity in both cell-attached voltage-clamp and whole-cell current-clamp recordings with intact synaptic transmission (Figure 1B-C, Chi squared test, p < 0.01). Moreover, we found that the percentage of neurons which were spontaneously active in tonic firing was larger in both types of recordings in adult male MD rats compared to control non-MD rats, while MD-induced increases in neuronal bursting were only detected in current-clamp recordings, similar to what we found in early and late adolescent male rats^5,7^.

**Figure 1.**
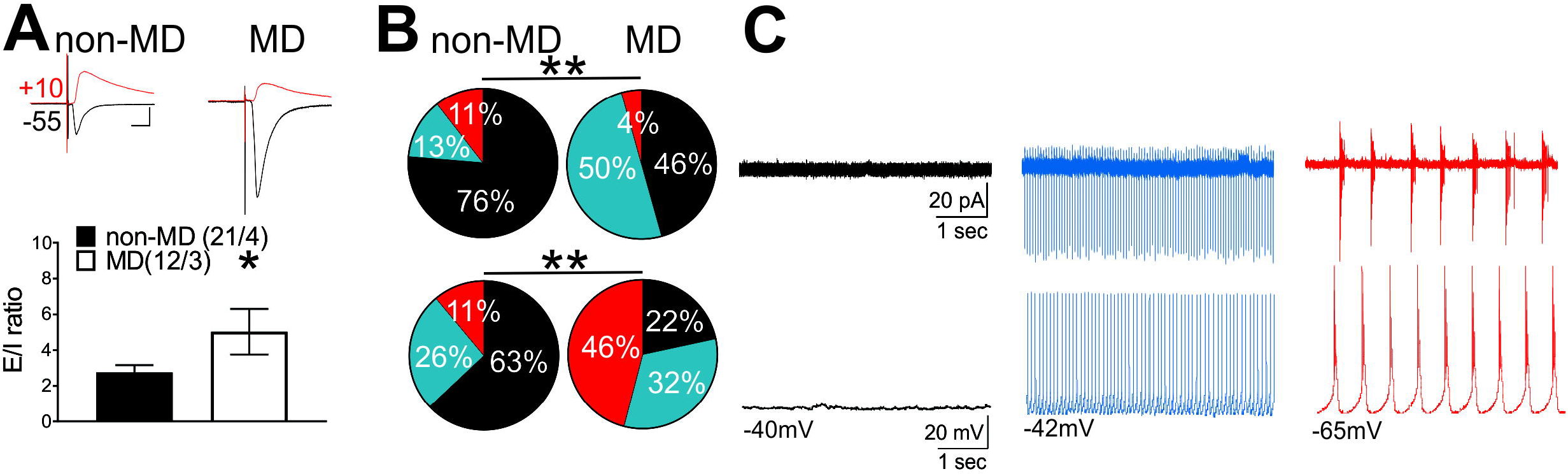
MD increased E/I ratios and spontaneous activity of LHb neurons. **A**, Summary of E/I ratios obtained from non-MD (n= 21/4) and MD rats (n=12/3), with representative traces of evoked EPSCs (black, recorded at −55-mV holding potential) and IPSCs (red, recorded at +10-mV holding potential) in response to the stria medullaris stimulation for LHb neurons. **B-C**, Pie charts with representative traces of voltage-clamp cell-attached recordings (top, V = 0, non-MD, n = 38/4; MD, n = 46/5) and current clamp whole-cell recordings (bottom, I = 0, non-MD, n = 27/4; MD, n = 37/5) of spontaneous neuronal activity across non-MD and MD rats. Comparison of the percent distributions of silent (black), tonic (blue), or bursting (red) LHb neurons showed a significant increase in tonic (both recordings) and bursting (only in current clamp recordings) LHb neuronal activity following MD. *p < 0.05, **p < 0.01 by unpaired Student’s t tests or Chi squared tests. In this and **Figure 3** n represents the number of recorded cells/rats.

### MD altered the acquisition of MSA and reduced sucrose preference in adulthood

We tested the effects of MD on intravenous MSA acquisition at 0.3 mg/kg/infusion or SSA on a FR1 schedule (morphine/saline delivered after one active lever press) for 6 days followed by a FR3 schedule (morphine/saline delivered after 3 active lever presses) of reinforcement for 4 days in adult male rats. We found that although both non-MD and MD rats readily acquired MSA compared to animals with SSA, MD rats significantly decreased the number of drug-paired lever presses as compared to non-MD rats in FR3 schedule sessions (Figure 2A; 3-way RM ANOVA tests; active lever presses with the FR1 schedule: effect of MD: F (1, 23) = 5.307, p<0.05; effect of MSA: F (1, 23) = 5.347, p<0.05; effect of time: F (5, 115) = 1.527, p=0.1867; MD × time interaction: F (5, 115) = 2.122, p=0.0677; MSA × time interaction: F (5, 115) = 0.9768, p=0.4351; MD × MSA interaction: F (1, 23) = 0.9037, p=0.3517; MD × MSA × time interaction: F (1, 23) = 0.9037, p<0.0001; active lever presses with the FR3 schedule: effect of MD: F (1, 23) = 8.371, p<0.01; effect of MSA: F (1, 23) = 43.07, p<0.0001; effect of time: F (3, 69) = 1.910, p=0.1360; MD × time interaction: F (3, 69) = 0.5974, p=0.6189; MSA × time interaction: F (3, 69) = 1.028, p=0.3857; MD × MSA interaction: F (1, 23) = 9.475, p<0.01; MD × MSA × time interaction: F (3, 69) = 0.6076, p=0.6123). MD rats also self-administered lower amount of morphine as compared to non-MD rats in FR3 schedule sessions (Figure 2B; 3-way RM ANOVA tests; morphine intake with the FR1 schedule: effect of MD: F (1, 23) = 3.107, p=0.0912; effect of MSA: F (1, 23) = 3.562, p=0.0718; effect of time: F (5, 115) = 2.303, p<0.05; MD × time interaction: F (5, 115) = 2.153, p=0.0642; MSA × time interaction: F (5, 115) = 5.053, p<0.001; MD × MSA interaction: F (1, 23) = 1.217, p=0.2813; MD × MSA × time interaction: F (5, 115) = 0.7924, p=0.5572; morphine intake with the FR3 schedule: effect of MD: F (1, 23) = 9.960, p<0.01; effect of MSA: F (1, 23) = 44.71, p<0.0001; effect of time: F (3, 69) = 0.9333, p=0.4294; MD × time interaction: F (3, 69) = 0.1069, p=0.9558; MSA × time interaction: F (3, 69) = 0.7312, p=0.5369; MD × MSA interaction: F (1, 23) = 10.01, p<0.01; MD × MSA × time interaction: F (3, 69) = 0.8669, p=0.4626). In both non-MD and MD rats, MSA increased locomotor activity in FR3 schedule sessions (Figure 2C, Mixed-effect analysis and 3-way RM ANOVA tests; locomotor activity with the FR1 schedule: effect of MD: F (1, 23) = 0.003450, p= 0.9537; effect of MSA: F (1, 23) = 11.56, p<0.01; effect of time: F (3.154, 65.60) = 2.673, p=0.0518; MD × time interaction: F (5, 104) = 0.9543, p=0.0677; MSA × time interaction: F (5, 104) = 2.638, p<0.05; MD × MSA interaction: F (1, 23) = 0.1469, p= 0.7051; MD × MSA × time interaction: F (5, 104) = 1.347, p= 0.2506; locomotor activity with the FR3 schedule: effect of MD: F (1, 23) = 0.2612, p=0.6142; effect of MSA: F (1, 23) = 30.67, p<0.0001; effect of time: F (2.163, 49.75) = 0.6315, P=0.5480; MD × time interaction: F (3, 69) = 0.3589, p=0.7829; MSA × time interaction: F (3, 69) = 1.209, p=0.3131; MD × MSA interaction: F (1, 23) = 0.4136, p=0.5265; MD × MSA × time interaction: F (3, 69) = 0.2856, p=0.8357). Note that there were no significant effects of MD or MSA on numbers of inactive lever presses among non-MD and MD rats in FR3 schedule sessions (Figure 2D; 3-way RM ANOVA tests; inactive lever presses with the FR1 schedule: effect of MD: F (1, 23) = 1.123, p=0.3003; effect of MSA: F (1, 23) = 4.395, p<0.05; effect of time: F (2.725, 62.68) = 6.090, p=0.0015; MD × time interaction: F (5, 115) = 1.039, p=0.3983; MSA × time interaction: F (5, 115) = 1.123, p=0.3521; MD × MSA interaction: F (1, 23) = 1.861, p=0.1857; MD × MSA × time interaction: F (5, 115) = 0.2369, p=0.9455; inactive lever presses with the FR3 schedule: effect of MD: F (1, 23) = 2.880, p=0.1032; effect of MSA: F (1, 23) = 0.8052, p=0.3788; effect of time: F (2.206, 50.74) = 0.1974, p=0.8417; MD × time interaction: F (3, 69) = 0.6188, p=0.6052; MSA × time interaction: F (3, 69) = 1.168, p=0.3283; MD × MSA interaction: F (1, 23) = 0.001577, p=0.9687; MD × MSA × time interaction: F (3, 69) = 0.4082, p=0.7476). We also tested the effects of MD on the SPT and found that MD rats exhibited a significantly lower sucrose consumption compared to non-MD rats, suggesting an anhedonia-like behavior in MD male rats (Figure 2E, t (17) = 2.35, p < 0.05).

**Figure 2.**
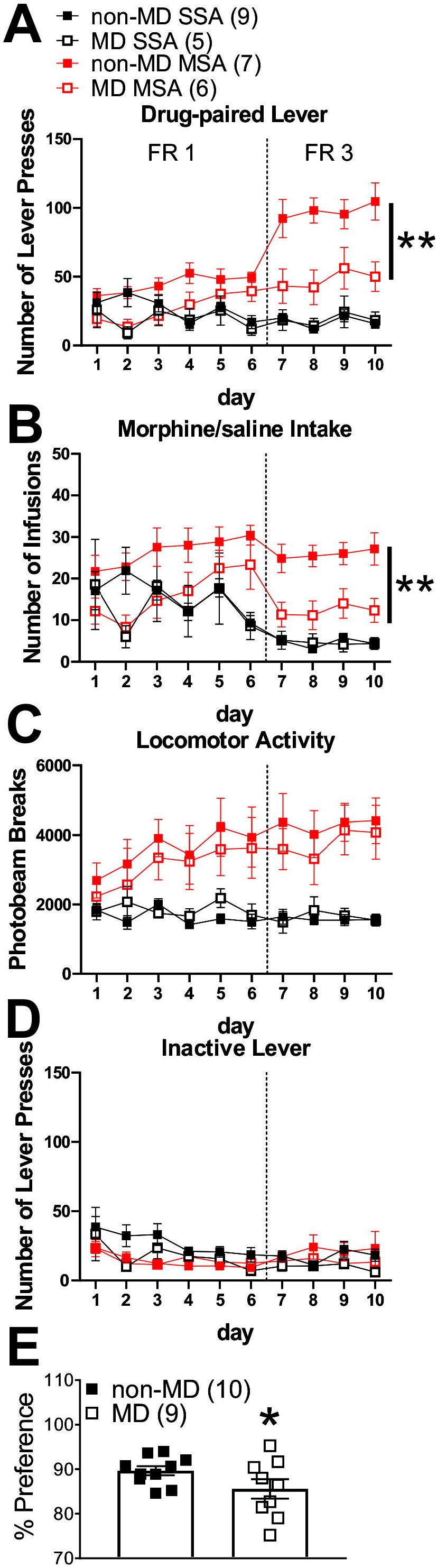
MD decreased morphine intake during FR3 MSA and sucrose preference in the SPT. Average number of active lever presses (**A**), drug infusions (**B**), locomotor activity (**C**) and inactive lever presses (**D**) during FR1 and FR3 SSA (black) and MSA (at 0.3 mg/kg per infusion of morphine, red) sessions. MD significantly decreased active lever presses and morphine intake in FR1 and FR3 sessions of MSA. In both non-MD and MD rats, MSA induced locomotor sensitization (n=5-9/group). **E**, MD animals consumed less sucrose solution compared to the control non-MD counterparts in the SPT (n=10/group). *p < 0.05, **p < 0.01 by 2-way ANOVA or unpaired Student’s t tests.

### MD and MSA potentiated glutamatergic synaptic transmission onto LHb neurons in adult male rats

We recorded AMPAR-mediated mEPSCs in non-MD and MD rats with SSA or MSA at 0.3mg/kg/infusion, one day after the last session of self-administration (Figure 3A-C). Similar to our observations in adolescent MD rats ^6^, glutamatergic synapses onto LHb neurons were potentiated both pre-and post-synaptically in adult MD rats with SSA compared to non-MD rats with SSA (effect of MD). The increase in postsynaptic AMPAR-mediated synaptic function was evident by a significant increase in the cumulative probability of AMPAR-mediated mEPSC amplitude after MD (Figure 3C, KS tests, p<0.0001), although the average amplitudes were not significantly different between MD and non-MD rats (Figure 3B, two-way ANOVA tests; effect of MD: F (1, 75) = 0.3144, p=0.5766; effect of MSA: F (1, 75) = 1.208, p=0.2752; MDXMSA interaction: F (1, 75) = 0.0005421, p=0.9815). Moreover, we detected a significant increase in the cumulative probability of AMPAR-mediated mEPSC frequency and the average frequencies in MD rats with SSA compared to those from non-MD rats with SSA, suggesting an increase in presynaptic glutamate release by MD (Figure 3B-C, KS tests p<0.0001; two-way ANOVA tests; effect of MD: F (1, 75) = 36.86, p<0.0001; effect of MSA: F (1, 75) = 6.577, p<0.01; MDXMSA interaction: F (1, 75) = 5.199, p<0.05). Similar pre- and post-synaptic glutamatergic plasticity was present in MD rats with MSA compared to non-MD SSA rats 24 h after the last morphine intake, suggesting that MD-induced glutamatergic plasticity that persists into adulthood remains intact in MD rats that self-administered morphine. Although it should be noted that there was a significant decrease in the mean and cumulative probability of mEPSC frequency in MD rats with MSA compared to MD rats with SSA (Figure 3, 2-way ANOVA and KS tests, p<0.01, p<0.0001). Interestingly, MSA also triggered a postsynaptic glutamatergic potentiation of LHb neurons in non-MD MSA rats compared to non-MD SSA rats (effect of MSA) (Figure 3, 2-way ANOVA and KS tests, p<0.01, p<0.0001). This was evident by a significant increase in the cumulative probability of AMPAR-mediated mEPSC amplitude after MSA (Figure 3C, KS tests, p<0.0001), although the average amplitudes were not significantly different between non-MD with SSA and non-MD with MSA rats (Figure 3B).

**Figure 3.**
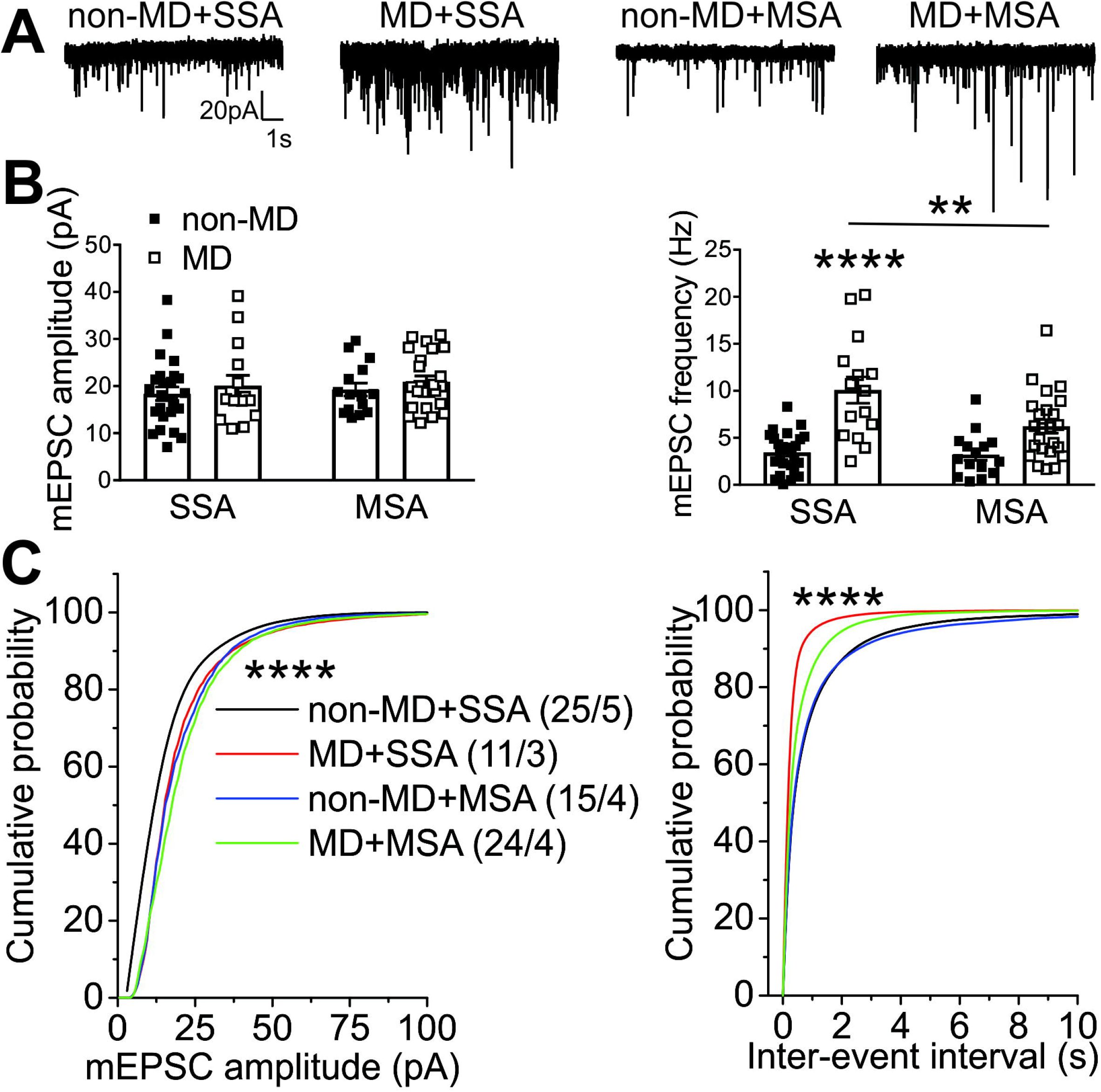
Effects of intravenous SSA or MSA 24h after the last morphine intake at LHb glutamatergic synapses from non-MD and MD rats. Representative AMPAR-mediated mEPSC traces from non-MD and MD rats (**A**, calibration bars, 20 pA/1 s), average mEPSC amplitude and frequency (**B**) and cumulative probability plots of amplitude and frequency (inter-event interval, **C**) in non-MD and MD rats (non-MD+SSA: n = 25/5, MD+SSA: n=11/3, non-MD+MSA: 15/4, MD+MSA: n=24/4). MSA and MD both potentiated glutamatergic synapses onto LHb neurons but this potentiation was also significantly decreased 24h following MSA in MD rats (**p<0.01, ****p<0.0001 by 2-Way ANOVA or Kolmogorov–Smirnov tests).

## Discussion

Although the general consensus in the literature is that ELS increases the risk of addiction, conflicting and sometimes opposite results have been reported in regards to drug seeking/taking behaviors. Previous work on the effects of ELS in drug addiction has largely focused on stimulants and alcohol ^12,15,30–40^ while a few studies report ELS effects in the context of opioid dependence and opioid reward using experimenter-driven passive morphine administration in morphine-induced sensitization and conditioned place preference (CPP)^23,41,42^. Here, we used intravenous MSA, which incorporates decision-making processes and reward circuitry to a greater extent than passive administration studies to assess how MD altered volitional morphine intake during MSA and LHb glutamatergic synaptic function following cessation of morphine intake 24h after the last MSA session in adult male rats. We provided evidence that MD-induced pre- and post-synaptic potentiation at glutamatergic synapses onto LHb neurons as well as increased E/I ratios demonstrating that LHb hyperactivity ^5–7^ persisted into adulthood in male MD rats. Consistent with our earlier observation of increased immobility in the FST in late adolescent MD rats ^5^, we found that adult MD rats also showed anhedonia-like behavior in the SPT, suggesting that a single prolonged MD stress promotes depressive- and anhedonic-like states in SD male rats. This is in agreement with other ELS studies using MD/MS procedures in SD rats that also trigger increased immobility in the FST and a loss of preference for sucrose as a natural reward ^43–45^. In contrast, an MS procedure that increased oral morphine self-administration behavior and preference in separated Long-Evans male rats showed that MS rats also enhanced their sucrose preference in the SPT ^23^. Regardless of the differences in rat strains and ELS models, it is notable that the SPT in this study was performed differently from the commonly used SPT in the literature. The SPT protocol in this study ^23^ lasted for 90 days and modified to measure preference to a very low concentration of sucrose (0.025% sucrose) that induced a preference of 70% in both control non-separated and separated rats as compared to higher concentrations of sucrose commonly used for the SPT including the 2% sucrose used in our MD study which also induced a high preference in their rats ^23^. Nevertheless, this study revealed that MS in Long-Evans male rats induced an initial decline (detected from days 11 to 34 which was also present in non-separated controls) and then a long-lasting increase (from days 47 to 54 and on day 90 only in separated rats) in sucrose preference ^23^ suggesting that MS stress may initially diminish but then accentuate the rewarding properties and perception of natural rewarding stimuli following long-term exposures. Interestingly, the same MS model increased morphine and amphetamine, but not cocaine and ethanol oral consumption and preference in Long-Evans male rats, suggesting that the ELS MS rat model is highly suitable to detect vulnerability to opioids^24^. In our study, both non-MD and MD rats self-administered morphine, suggesting that rats perceived morphine as rewarding compared to saline administration. Although, morphine induced locomotor sensitization during MSA acquisition in both non-MD and MD rats, MD rats significantly took less morphine compared to non-MD rats. Given that LHb hyperactivity and glutamatergic potentiation in LHb neurons underlie negative affective disorders including depression and drug withdrawal ^46^, we also predicted that the lower morphine intake in MSA may be associated with LHb glutamatergic dysfunction that could mediate not only MD-induced sucrose anhedonia but also highlight aversive properties of morphine in MD rats. Similar to our observations in adolescent MD rats^6^, glutamatergic synapses onto LHb neurons were still potentiated pre-and post-synaptically in adult MD rats self-administering saline. The postsynaptic glutamatergic plasticity remained intact in LHb of MD rats with MSA 24h following their last morphine intake and although morphine was able to significantly decrease part of the MD-induced presynaptic potentiation, the increased presynaptic glutamate remained significantly higher compared to control non-MD rats self-administering saline. Interestingly, we also detected an induction of postsynaptic glutamatergic potentiation in LHb neurons of non-MD MSA rats compared to non-MD SSA rats 24h following the last MSA. We assume that LHb glutamatergic potentiation in control non-MD rats may signal the absence of expected morphine reward 24h after their last infusion while in MD rats, this persistent plasticity may mediate aversive properties of morphine that limited morphine intake in MD rats. Similarly, LHb activity is linked to aversive properties of cocaine. While passive injections of cocaine in mice have been shown to potentiate AMPAR-mediated glutamatergic plasticity in LHb neurons ^47–49^, cocaine exhibits a bi-phasic response in LHb with an initial inhibition followed by delayed excitation of a subset of LHb neurons in rats which mediates cocaine’s initial rewarding effects and delayed aversive properties ^50^. On the other hand, a recent study showed that naloxone-precipitated withdrawal from passive and repeated administration of morphine in control mice triggers depression rather than potentiation at glutamatergic synapses onto LHb neurons projecting to raphe nucleus ^27^, suggesting the complexity of drug modulation of specific LHb neuronal circuits. While it is possible that morphine also induces biphasic activation of a specific subpopulation of LHb neurons similar to cocaine, it is important to note that these studies implemented an experimenter-administered cocaine or morphine rather than volitional drug intake used in our MSA rat studies where synaptic mechanisms underlying reward learning and decision-making processes in reward circuits are engaged to a greater extent than passive administration.

Studies using cocaine self-administration also indicate the involvement of LHb activity in mediating the negative outcomes of drug-seeking behaviors ^51,52^. For example, in the ELS model of limited bedding and nesting (LBN, a naturalistic rodent model of ELS), LBN rats self-administer lower unit doses of cocaine compared to controls possibly due to a persistent anhedonia induced by LBN and altered activity in reward and stress related areas including LHb 12,35. In general, withdrawal from passive repeated in vivo ethanol administration or voluntary ethanol consumption under an intermittent access two-bottle choice procedure in rats promote depressive and anxiety-like behaviors and also increases LHb spontaneous activity and potentiates glutamatergic synapses in LHb ^53,54^. Ethanol withdrawal-induced decreased expression of LHb astrocyte glial glutamate transporter (GLT-1) that limits the uptake of synaptic glutamate underlies LHb glutamatergic potentiation and hyperactivity as well as depression and anxiety during ethanol withdrawal ^54^. It would be interesting to test whether MD-induced glutamatergic potentiation and hyperactivity involve dysfunctional astrocytic GLT-1 function which could also mediate anhedonic- and depressive-like behaviors as well as contribute to aversive properties of morphine and reduced morphine intake during MSA in MD rats.

Although we have favored the idea that MD-induced LHb hyperactivity reduces rewarding properties of morphine and mediates anhedonic and aversive aspects of opioid intake in MD rats, it is also possible that LHb hyperactivity associated with MD could result in increased sensitivity to low doses of morphine. This warrants investigating the effects of MD on a morphine dose-response curve for MSA given that MD also increased locomotor sensitization to morphine. Of note, it has been documented that brief MS (e.g., 15-min daily separations of the litter from the dam, often referred to as “handling,”) may reduce the rewarding effects of drugs of abuse while longer MS/MD procedures may enhance the reinforcing properties of drugs including opioids in adulthood ^23,24,36,41,55,56^. Thus, differences between predictable (repeated MS and MD) and unpredictable (our single prolonged MD and LBN) stressors as well as the duration of MD/MS in ELS models may confer resistance or vulnerability to drugs of abuse and directly impact the outcomes in terms of addictive behaviors as well as comorbid depression/mood phenotypes. Given that our MD rat model is a single prolonged and unpredictable early life stressor and is associated with depressive- and anhedonia-like behaviors as well as changes in MSA acquisition, it may represent a valid ELS model for investigation of potential therapeutic interventions for treatment of opioid use disorder and comorbid mood disorders in patients with a history of childhood neglect and abuse. Our recent discoveries of the neuromodulatory regulation of LHb neuronal excitability and synaptic transmission by intra-LHb corticotropin-releasing factor and dynorphin/Kappa opioid receptor signaling and their dysregulation by MD ^6,7^ further highlight the possible involvements of some of the critical and less-studied synaptic and molecular mechanisms underlying ELS-induced neuromodulation within LHb circuits that could affect the rewarding and negative reinforcing properties of opioids and promote opioid seeking behaviors.

## Acknowledgments

The opinions and assertions contained herein are the private opinions of the authors and are not to be construed as official or reflecting the views of the Uniformed Services University of the Health Sciences or the Department of Defense or the Government of the United States. This work was supported by the National Institute of Drugs of Abuse (NIH/NIDA) Grant#R01 DA039533 to FN. The funding agency did not contribute to writing this article or deciding to submit it. We would like to thank Dr. Brian Cox for his helpful and critical review of the manuscript and the support of the Rat Behavior Core at USU.

## Author Contribution

FN, LL and KC were responsible for the study concept and design. LL, RB, RS, SS, MT, and SG contributed to the acquisition of animal data. FN, LL, RB, RS, SS, MT, SG and KC assisted with data analysis and interpretation of findings. FN, LL and KC wrote the manuscript. All authors critically reviewed content and approved final version of manuscript for submission.

## Data Sharing

The data that support the findings of this study are available on request from the corresponding author. The data are not publicly available due to privacy or ethical restrictions.

